# Constitutive cGAS-STING activation in ALT+ cells

**DOI:** 10.1101/2025.04.18.649516

**Authors:** Dan Geelen, Alexander Beacom, Mélina Vaurs, Manon Mahieu, Rachid Boutoual, Amélie Derumier, Remington Hoerr, Axelle Loriot, Joe Nassour, Anabelle Decottignies

## Abstract

Stimulator of interferon genes (STING) is a pivotal mediator of anti-tumor immunity, activated downstream of cytoplasmic DNA recognition by cyclic GMP-AMP synthase (cGAS). In cells that employ the alternative lengthening of telomeres (ALT) pathway, extrachromosomal telomeric repeats (ECTRs) act as potent activators of the cGAS–STING pathway. Although ALT+ cells were previously thought to evade this pathway through epigenetic silencing, our findings reveal more nuanced adaptive mechanisms that involve the spatial regulation of STING and the clearance of immunostimulatory ECTR species. In the vast majority of ALT+ cells, ongoing ECTR production drives sustained 2′3′-cGAMP synthesis, directing STING to the Golgi apparatus for immune signaling before lysosomal degradation. Lysosomal degradation of STING relies largely on interferon regulatory factor 3 (IRF3), with minimal involvement of TANK-binding kinase 1 (TBK1), pointing at a novel role for IRF3 in STING trafficking in ALT+ cells. Another adaptive mechanism involves the removal of immunogenic cytoplasmic ECTRs through STING-mediated lysosomal degradation. These mechanisms likely cooperate to restrict chronic interferon signaling that might otherwise prevent ALT+ cancers from emerging. These findings refine our understanding of immune sensing in ALT+ cancers, revealing how ALT+ cells respond to continuous ECTR production and suggesting a potential therapeutic target to modulate the microenvironment of ALT+ tumors.

## Introduction

Stimulator of interferon genes (STING) serves as a powerful inducer of anti-tumor immunity (Corrales et al., 2015 and 2016). STING engagement can occur downstream of cytoplasmic DNA recognition by cyclic guanosine monophosphate (GMP)-AMP (cGAMP) synthetase (cGAS). Cytoplasmic cGAS ligands include pathogenic DNA from bacteria or viruses, as well as self-DNA fragments, such as those arising from chromosomal instability or released from the mitochondria (Kim et al., 2023; Bakhoum and Cantley, 2018; Beernaert and Parkes, 2023). The immune functions of STING are highly dependent on intracellular trafficking (Balka and De Nardo, 2021). When cGAS detects cytoplasmic DNA, it produces 2’3’-cGAMP, which subsequently binds to the endoplasmic reticulum (ER)-localized adaptor STING, triggering its conformational change and relocalization to the ER-Golgi intermediate compartment (ERGIC), and then to the Golgi (Wu et al., 2013). At the trans-Golgi network, STING associates with the downstream kinase TANK-binding kinase 1 (TBK1), triggering its activation and subsequently leading to the transcription of target genes via interferon regulatory factor 3 (IRF3) (Chauvin et al., 2023). STING activation can also induce nuclear factor (NF)-κB signaling, although less is known about the precise mechanism of this response. The induction of IRF3 and NF-κB signaling leads to a transcriptional response that includes type I interferons (IFNs) and interferon-stimulated genes (ISGs), leading to the production of proinflammatory cytokines and chemokines (Decout et al., 2021).

Multiple processes at various stages take place to terminate STING signaling to prevent the over activation of the immune response. First, STING protein undergoes various post-translational modifications, such as phosphorylation and ubiquitination, that promote its trafficking to the endolysosomal compartment for degradation (Gentili et al., 2023; Kuchitsu et al., 2023; Balka et al., 2023; Zhao et al., 2024; Liu et al., 2022; Xu and Wan, 2024). In a similar manner, ubiquitinated IRF3 is subjected to autophagic degradation, which further aids in terminating the type I IFN response (Zhang et al., 2008). A third mechanism of termination involves the clearance of cytoplasmic DNA through a mechanism distinct from conventional autophagy and occurring independently of interferon induction (Gui et al., 2019). This subsequent termination mechanism is mediated by 2’3’cGAMP-bound STING at the ERGIC, which ultimately guides the cytosolic DNA molecules toward lysosomal degradation (Gui et al., 2019).

Despite cGAS-STING pathway’s canonical role as a potent activator of anti-tumor immunity, growing evidence, including the role of cGAS-STING in clearing cytoplasm DNA, points, paradoxically, to its contribution to cancer cell survival. In fact, this pathway remains active in certain tumor cells with chromosomal instability (Corrales et al., 2015 and 2016; Sivick et al., 2018; Beernaert and Parkes, 2023). In addition, in cancer cells with elevated chromosomal instability, cGAS-STING activation fosters pro-tumoral activity that depends on NF-κB and IL-6 (Bakhoum et al., 2018; Hong et al., 2022; Lanng et al., 2024). This process is mediated through the activation of extracellular signal-regulated kinase 1/2 (ERK1/2), a key downstream effector of the IL-6/JAK signaling pathway, which plays a key role in cancer development by regulating essential cellular processes such as proliferation, migration, and stress responses (Guo et al., 2020).

Alternative lengthening of telomeres (ALT) constitutes a homology-directed repair mechanism that a subset of cancers employs to sustain telomere length in the absence of telomerase. A defining characteristic of ALT-positive (ALT+) cells is the production of abundant extrachromosomal telomeric repeats (ECTRs), which can potentially activate cytosolic DNA-sensing pathways such as cGAS–STING and drive the transcription of ISGs (Chen et al., 2017; Nassour et al., 2019, 2023, 2024; Siametis et al., 2024). Previous work suggested that ALT development involves the silencing of the *STING* gene as a strategy to evade the activation of the DNA sensing pathway triggered by ECTRs, accomplished through (or as a byproduct of) loss of functional ATRX-DAXX histone chaperone complex (Chen et al., 2017). While this work supported the potential incompatibility of a functional cGAS-STING pathway with the ALT+ phenotype, other studies demonstrated that ALT+ cancer cells could activate the cGAS/STING pathway in response to either the accumulation of cytoplasmic ECTRs following chromatin decompaction (Segura-Bayona et al., 2020) or the degradation of stalled replication forks (Emam et al., 2022). Furthermore, the degradation of cytoplasmic telomeric DNA fragments through cGAS-dependent autophagy was recently described as a survival mechanism in RAD51AP1-depleted ALT+ cells experiencing telomere dysfunction (Barroso-Gonzales et al., 2019). Although these later studies showed that the cGAS-STING pathway could be activated in ALT+ cells, it remained unclear whether the naturally occurring cytoplasmic ECTRs in ALT+ cells drive constitutive activation of the cGAS-STING pathway.

Here, we found that, while some ALT+ cell lines indeed repress pathway activity through *cGAS* or *STING* transcriptional downregulation, most keep the expression of both genes and exhibit pronounced pathway activation. In the cell lines with functional cGAS-STING signaling, the continuous leakage of ECTRs to the cytoplasm stimulates strong cGAS activity, resulting in increased 2’3’-cGAMP synthesis and triggering STING translocation from the ER to the Golgi apparatus. At the Golgi, STING promotes innate immune signaling, after which it undergoes lysosomal degradation. Unexpectedly, this lysosomal degradation relies upon the presence of IRF3, with loss of IRF3 protein completely recovering STING protein levels. In addition to promoting inflammatory signaling, STING facilitates ECTR clearance from the cytoplasm through lysosomal degradation. Collectively, these observations refine the prevailing model of immune sensing in ALT+ cancers.

## Results

### Persistent lysosomal degradation of STING in ALT+ cells

To examine cGAS-STING regulation in ALT+ cancers, we built a panel of sixteen human cell lines, including nine ALT+ cell lines, four telomerase-positive (TEL+) cell lines, and three primary human fibroblast lines (IMR90, WI38, MRC5). Four cell lines were derived from patients with osteosarcomas (U2OS/ALT+, SaOS-2/ALT+, G292/ALT+, HOS/TEL+), four from patients with adenocarcinomas (TOV-112D/ALT+, Calu-3/TEL+, HeLa-LT/TEL+, HT-29/TEL+), and one from a patient with a rhabdomyosarcoma (SJCRH30/ALT+) or neuroblastoma (SK-N-F1/ALT+). The panel also included three ALT+ *in vitro* immortalized fibroblast lines (GM847/ALT+, WI38-VA13/ALT+, KMST-6/ALT+). Consistent with the previously described frequent loss of ATRX or DAXX in ALT+ cell lines (Lovejoy et al., 2012), only two ALT+ cell lines (TOV-112D and SJCRH30) express normal levels of both full-length ATRX and DAXX (Fig S1A), although their functionality has not been assessed.

We first characterized the amount of cGAS and STING protein expressed in each cell line by immunoblot. All the TEL+ and primary cell lines expressed readily detectable amounts of both cGAS and STING (Fig 1A and FigS1B). However, in agreement with previous work (Chen et al., 2017), SK-N-F1 and TOV-112D ALT+ cell lines lacked detectable cGAS protein, and eight of the nine ALT+ cell lines expressed weak or undetectable amounts of STING protein (Fig 1A and Fig S1B). In these cell lines, the levels of *STING* mRNA were highly variable and did not correlate with the STING protein expression. For instance, cell lines like SJCRH30 and WI38-VA13 displayed high *STING* mRNA levels by RT-qPCR but had barely detectable protein amounts by immunoblot, suggesting post-translational regulation. Conversely, in KMST-6 cells with undetectable *cGAS* mRNA expression, both *STING* mRNA and protein levels were high (Fig 1A and Fig S1B-C).

**Figure 1.**
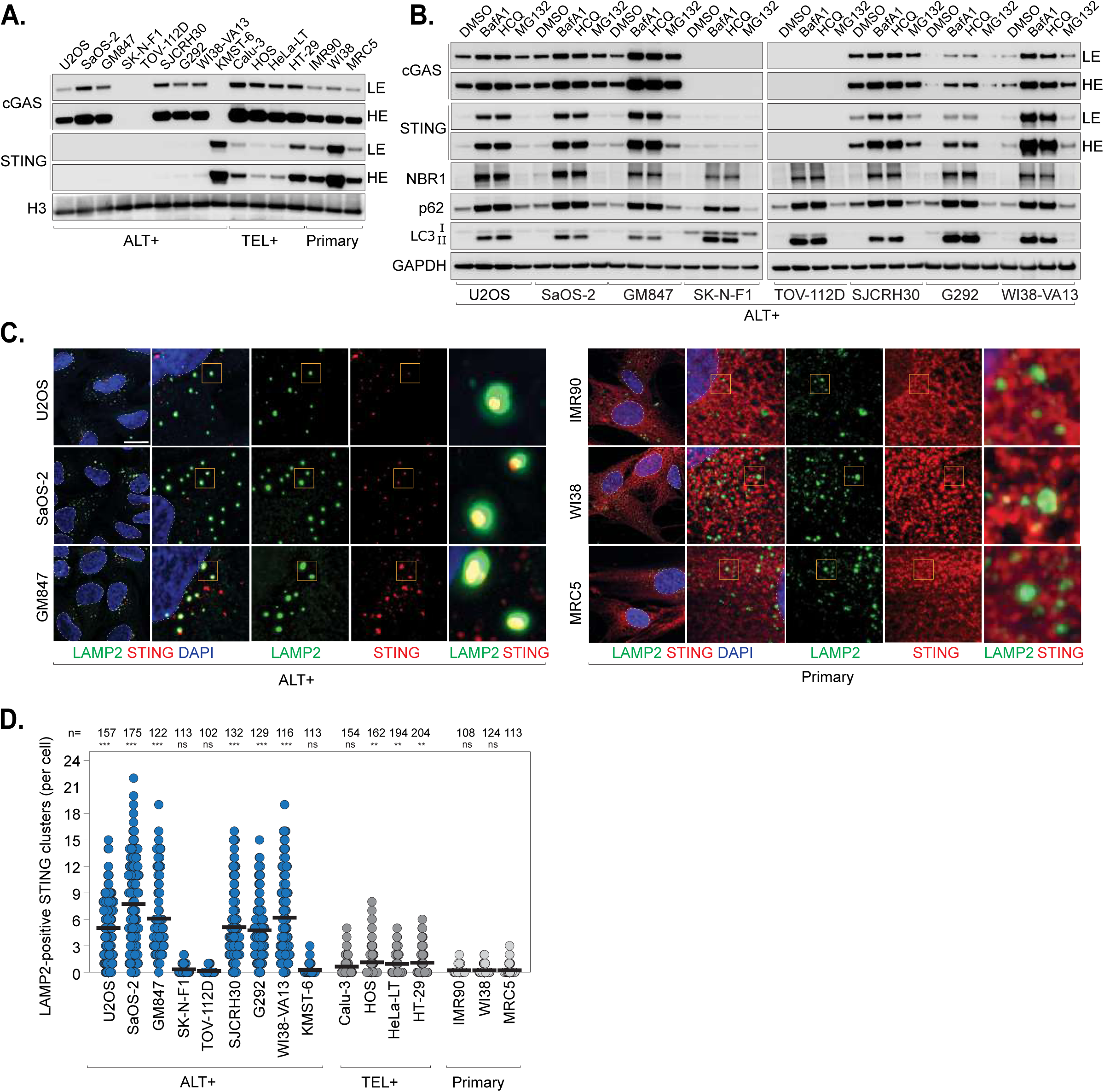
Persistent lysosomal degradation of STING in ALT+ cells. **(A)** Immunoblot analysis of cGAS and STING in primary human fibroblasts (IMR90, WI38, MRC5), ALT+ (U2OS, SaOS-2, GM847, SK-N-F1, TOV-112D, SJCRH30, G292, WI38-VA13, KMST-6) and TEL+ cell lines (Calu-3, HOS, HeLa-LT, HT-29). Histone H3 (H3) was used as a loading control. n=3. LE (low exposure), HE (high exposure). Quantification is shown in Fig S1B. **(B)** Immunoblot analysis of the indicated ALT+ cells treated with either DMSO (control), the lysosomal inhibitors bafilomycin A1 (BafA1) or hydroxychloroquine (HCQ), or the proteasome inhibitor MG132 for 24 h. n=3. GAPDH was used as a loading control. RT-qPCR analysis of *STING* mRNA levels in the corresponding samples is shown in Fig S1D. **(C)** Apotome maximum intensity projection images of the indicated cells, co-immunostained with anti-LAMP2 and anti-STING. n=3. Scale bar, 20 μm. **(D)** Number of STING cytosolic clusters colocalizing with the lysosomal marker LAMP2. Mean. n=3, with the total number of cells counted indicated above. Kruskal–Wallis test: samples were compared individually to primary MRC5 cells. ns = not significant; \**p* < 0.05; \*\**p* < 0.01; \*\*\**p* < 0.001.

Lysosomal control of cGAS-STING signaling was characterized only recently (Xu and Wan, 2024), leading us to wonder if the lack of observable STING protein was due to lysosomal degradation. To test this, we treated the eight STING-negative cell lines with the lysosomal inhibitors bafilomycin A1 (BafA1) or hydroxychloroquine (HCQ), using the MG132 proteasome inhibitor as control (Rock et al., 1994; Yamamoto et al., 1998; Boya et al., 2005). Treatment with BafA1 or HCQ resulted in the accumulation of the known substrates of lysosomal degradation NBR1, p62, and LC3 in all cell lines, confirming treatment efficacy (Kirkin et al., 2009; Bjørkøy et al., 2005; Kabeya et al., 2000) (Fig 1B). Lysosomal inhibition, but not proteasome inhibition, led to a strong recovery of STING protein in all cell lines, except for SK-N-F1 and TOV-112D ALT+ cell lines, which lack cGAS protein (Fig 1B and Fig S1B). Treatment with lysosomal inhibitors also elevated *STING* mRNA levels (Fig S1D), consistent with the fact that STING itself functions as an ISG that promotes its own expression (Ma et al., 2015).

In support of the model suggesting active degradation of STING protein via the lysosome in most ALT+ cell lines, immunofluorescence imaging reveals weak cytoplasmic STING staining accompanied by variable numbers of clusters that colocalize with LAMP2, a key lysosomal membrane marker (Qiao et al., 2023), in these cell lines (Fig 1C-D). On the other hand, TEL+ or primary cell lines revealed strong cytosolic staining of STING that did not colocalize with LAMP2 (Fig 1C-D). Consistent with this, staining for calnexin, a molecular chaperone commonly used as a marker for the ER (Paskevicius et al., 2023), revealed colocalization of STING with calnexin in nearly all primary cells and about half of the TEL+ cells. However, in the ALT+ cells, only the cGAS-negative cell lines showed more than 10% of cells with STING localized to the ER (Fig S1E-F).

These data show that despite minimal observable STING protein in eight of the nine ALT+ cancer cell lines tested, the *STING* gene is generally not silenced. While STING localizes to the ER in its resting state in TEL+ and primary cells, in ALT+ cells, STING primarily localizes to the lysosome, indicating STING activation and trafficking. Inhibition of lysosomal degradation results in strong recovery of STING in ALT+ cells expressing cGAS, demonstrating that decreased protein levels observed in ALT+ cells are due to constitutive lysosomal degradation.

### cGAS–IRF3-dependent regulation of STING turnover in ALT+ cells

The observation of STING localization to the lysosome in ALT+ cells supports an active cGAS-STING pathway. Since STING must be activated by cyclic dinucleotide binding before exiting the ER, trafficking to the Golgi, and being delivered to the lysosome (Wu et al., 2013), we aimed to investigate whether interfering with pathway activation would prevent the lysosomal degradation of STING in ALT+ cells. To test this, we used CRISPR to selectively knock out cGAS in the STING-negative cell lines that express the gene. Accordingly, the loss of cGAS restored STING protein levels in all cell lines (Fig 2A). These findings are consistent with the observation of high STING protein levels in the cGAS-negative KMST-6 cells (Fig 1A). We next tested whether downstream actors of the cGAS-STING pathway, such as IRF3 and TBK1, may similarly impact STING protein levels. Surprisingly, loss of IRF3 led to a strong recovery of STING, while TBK1 loss had minimal effect on STING protein levels. With an alternative approach using siRNA against the same targets, we confirmed the degradation of STING in ALT+ cancer cells depends on cGAS and IRF3 (Fig S2). As phosphorylation of IRF3 by TBK1 is required for IRF3’s canonical role in promoting transcription of ISGs, these experiments have uncovered a novel, likely TBK1-independent, role of IRF3 in the degradation of STING in ALT+ cells (Tanaka and Chen, 2012) (Fig 2A).

**Figure 2.**
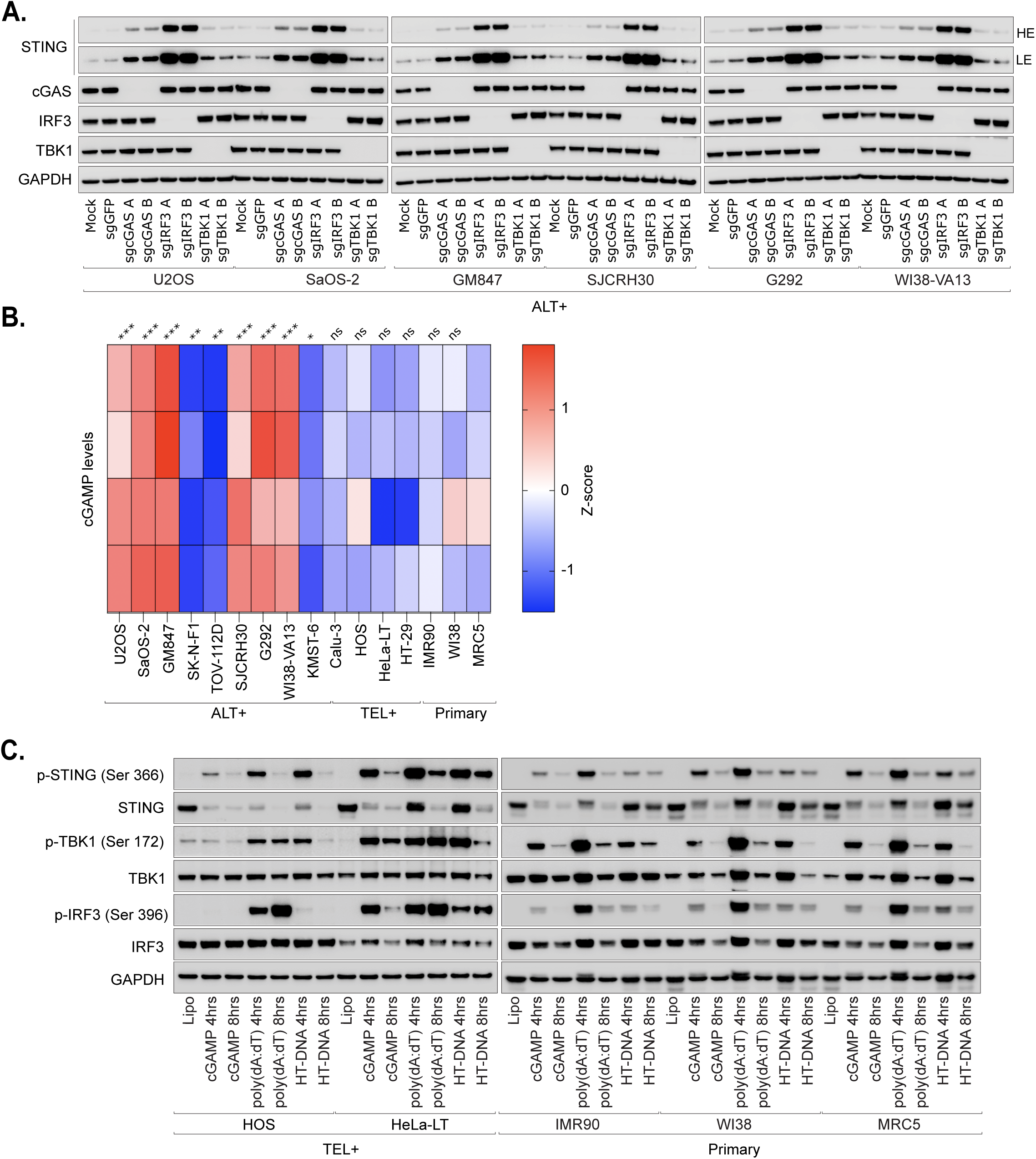
cGAS–IRF3-dependent regulation of STING turnover in ALT+ cells. **(A)** Immunoblot analysis of the indicated ALT+ cells transduced with lenti-CRISPR targeting cGAS, IRF3, or TBK1. Two distinct gRNAs were used for each gene. Cells transduced with a gRNA targeting GFP served as a control. Protein extracts were collected on day 6 post-transduction. GAPDH was used as a loading control. LE (low exposure), HE (high exposure). **(B)** 2′3′-cGAMP levels measured by ELISA in protein extracts from ALT+, TEL+ or primary cells, as indicated. Equal amounts of protein were used. n=4. 2’3’-cGAMP levels were log2 normalized and z-score values were calculated for each biological replicate. One-way ANOVA: samples were compared individually to primary MRC5 cells. ns = not significant; \**p* < 0.05; \*\**p* < 0.01; \*\*\**p* < 0.001. **(C)** Immunoblot analysis of the indicated TEL+ or primary cells treated with 2′3′-cGAMP (10 μg/mL) or transfected with poly(dA:dT) (2 μg/mL) or High-molecular-weight DNA (HT-DNA) (2 μg/mL). Protein extracts were collected at 4 h and 8 h post-treatment or post-transfection as indicated. Lipofectamine alone (Lipo) served as a control. GAPDH was used as a loading control.

As STING degradation in ALT+ cells was dependent on the presence of cGAS, we then investigated if cGAS enzymatic activity was increased in these cells. Using an ELISA to measure intracellular 2’3’-cGAMP, we determined all ALT+ cell lines, except for those lacking cGAS, contained significantly greater amounts of 2’3’-cGAMP than the TEL+ or primary cell lines (Fig 2B). Treatment of TEL+ or primary cells with 2’3’-cGAMP was sufficient to induce STING degradation, suggesting that the lack of constitutive STING degradation in these cells was not due to pathway dysfunction (Fig 2C). Similarly, poly(dA:dT) or herring testes (HT) DNA consistently resulted in phosphorylation of STING, TBK1, and IRF3 and subsequent degradation of STING in TEL+ or primary cells (Fig 2C).

Taken together, these data show that cGAS is highly active in ALT+ cells, producing significantly more intracellular 2’3’-cGAMP than in TEL+ or primary cells. Furthermore, the lysosomal degradation of STING is dependent on both cGAS and IRF3, as loss of either protein is sufficient to abolish STING degradation.

### Active cGAS-STING pathway in ALT+ cells

The continuous lysosomal degradation of STING protein in ALT+ cells, driven by cGAS, along with elevated 2’3’-cGAMP levels, indicated chronic activation of the pathway. To isolate the effect of the telomere maintenance mechanism on cGAS-STING pathway activity, we measured the expression of ISG transcripts in ALT+ and TEL+ cell lines with similar *cGAS*/*STING* mRNA levels and genetic backgrounds. For this purpose, we utilized the previously described TEL+ (n=3) and ALT+ (n=3) hybrid cell lines, which were derived from the *in vitro* immortalization of IMR90 human fibroblasts using SV40T (Episkopou et al., 2014 and 2019; Raghunandan et al., 2021). All ALT+ hybrids lack ATRX expression (Episkopou et al., 2019), and the mRNA levels of *cGAS* and *STING* were consistent across the hybrids (Fig S3A). Supporting our hypothesis, a GSEA analysis of RNA sequencing data from ALT+ and TEL+ hybrids showed that interferon-stimulated genes (ISGs) were significantly enriched among the genes upregulated in ALT+ cells (p=4.3 x 10^-26^), a finding we validated through RT-qPCR (Fig 3A and Fig S3B). The observed reduction in the mRNA levels of ISGs upon cGAS or STING depletion in SI24/ALT+ hybrid cells confirmed the functionality of the pathway in ALT+ cells (Fig 3B). The effect of cGAS knockdown was less pronounced compared to STING depletion, possibly due to the reduced cGAS knockdown efficiency.

**Figure 3.**
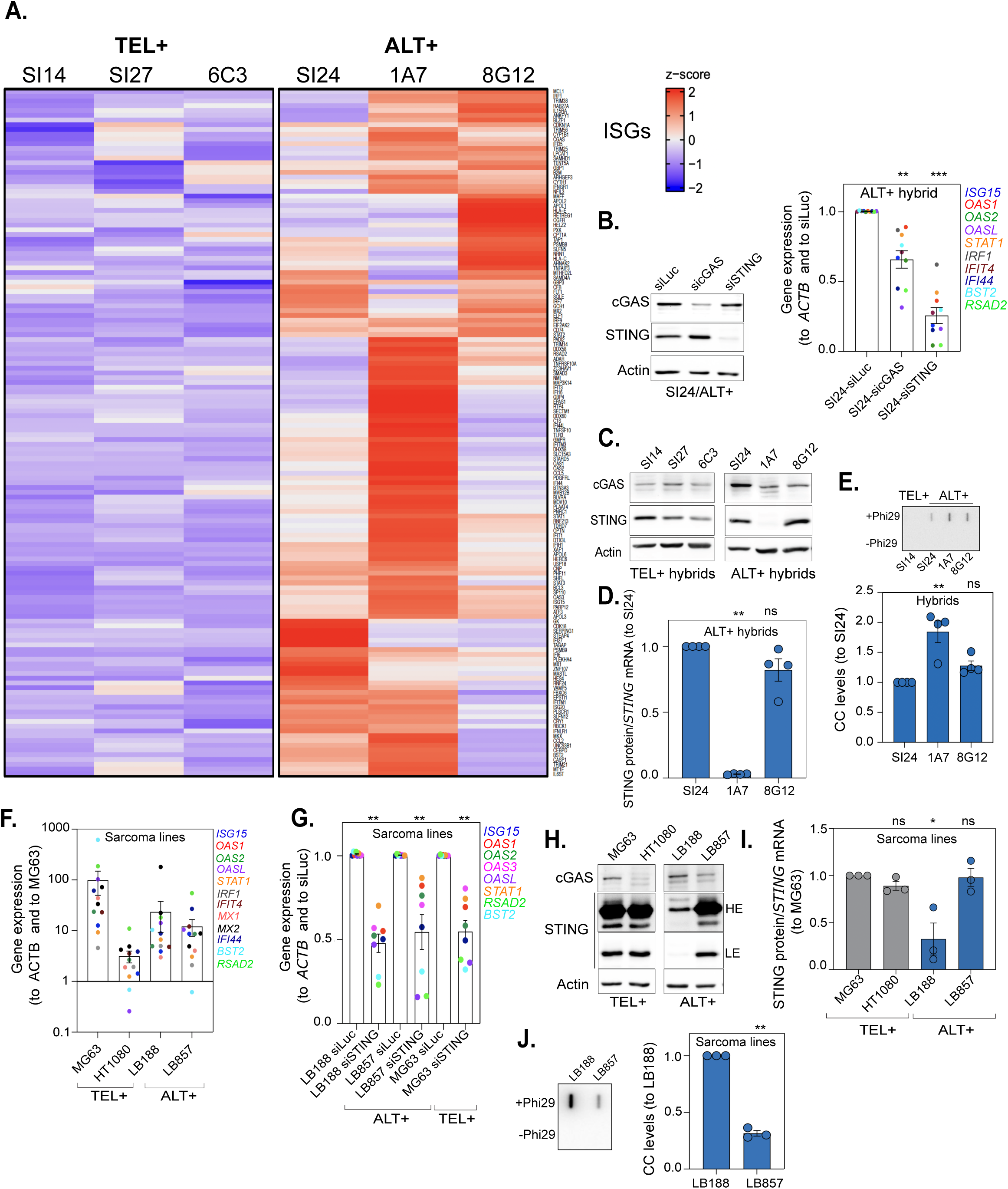
Constitutive activation of a functional cGAS-STING pathway in ALT+ cells. **(A)** Heatmap displaying the RNA-seq expression levels of ISG core-enrichment genes, accounting for the GSEA enrichment signal in ALT+ vs TEL+ hybrids. One experiment was performed. **(B)** Left: representative immunoblot analysis for knock-down control. Right: RT-qPCR analysis of the indicated ISGs upon cGAS or STING knock-down in SI24/ALT+ hybrid cells. Mean ± SEM. n=3-4. Ratio paired *t* tests: samples were compared individually to SI24-siLuc. **(C)** Representative immunoblot analysis of cGAS and STING in the hybrid cell lines. Actin was used as a loading control. n=4. **(D)** STING protein/mean *STING* mRNA levels shown for the indicated ALT+ hybrids were obtained from (C) and from Fig S3A, and normalized to SI24. Mean ± SEM. n=4. Kruskal-Wallis test: samples were compared individually to SI24. **(E)** Above: C-circle assays in the indicated ALT+ hybrids. Controls without Phi29 are shown. Below: quantification. Mean ± SEM. n=3. Kruskal-Wallis test: samples were compared individually to SI24. **(F)** RT-qPCR analysis of ISGs in the indicated sarcoma cell lines. Mean ± SEM. n=2-3. **(G)** RT-qPCR analysis of selected ISG genes upon STING knock-down in the indicated sarcoma cell lines. Cells were treated with siSTING for 72h; siLuc was used as control. Mean ± SEM. n=2-3. Ratio paired *t* tests. Validation of STING knock-down is shown in Fig S3F. **(H)** Representative immunoblot analysis of cGAS and STING in sarcoma cell lines, using Actin as loading control. n=3. LE (low exposure), HE (high exposure). **(I)** Same as (D) in the sarcoma cell lines, using data shown in (H) and in Fig S3E. Mean ± SEM. n=3. Kruskal-Wallis tests: samples were compared individually to MG63. **(J)** Same as (E) in LB188/ALT+ and LB857/ALT+ sarcoma cell lines. Mean ± SEM. n=3. Ratio paired *t* test. For all panels: ns = not significant; \**p* < 0.05; \*\**p* < 0.01; \*\*\**p* < 0.001.

Notably, among the ALT+ hybrids, the 1A7/ALT+ cell line showed the highest ISG expression levels (Fig 3A and Fig S3B). The 1A7/ALT+ cells exhibited a consistently lower STING protein-to-transcript ratio compared to the SI24/ALT+ and 8G12/ALT+ cell lines (Fig 3C-D and Fig S3A), with STING protein levels being restored upon treatment with BafA1 (Fig S3C), supporting the link between pathway activation and lysosomal degradation of STING. Assessing the LC3-II/LC3-I ratio, an indicator of autophagic flux, indicated that macroautophagy was, however, not generally elevated in 1A7/ALT+ cells (Fig S3D). Given that ECTRs have been previously linked to cGAS-STING pathway activation (Chen et al., 2017; Nassour et al., 2019, 2023, 2024; Siametis et al., 2024), we hypothesized that the 1A7/ALT+ cells might produce elevated levels of ECTRs, potentially explaining the increased degradation of STING protein. To test this, we measured ECTR levels using the established C-circle (CC) assay (Henson et al., 2017). Accordingly, CC levels were higher in 1A7/ALT+ cells (Fig 3E).

Next, to evaluate if the increased CC production and ISG expression found in the ALT+ hybrids also occurred in ALT+ cancers, we first screened a panel of eight TEL+ and ALT+ sarcoma cell lines for comparable *cGAS*/*STING* mRNA levels. Two TEL+ cell lines, MG63 (osteosarcoma) and HT1080 (fibrosarcoma), and two ALT+ cell lines, LB188 (rhabdomyosarcoma) and LB857 (myxoid sarcoma) were selected (Fig S3E). In contrast to the findings in hybrid cell lines, however, ISG expression was not consistently upregulated in the ALT+ sarcoma cell lines, and ISG transcript levels were higher in MG63/TEL+ cells (Fig 3F). This suggests that additional sources of self-DNA contribute to the activation of the cGAS-STING pathway in some cancer cells. In this context, mtDNA release, endogenous retroelements, chromatin alterations, chromosomal instability, and micronuclei—all potent activators of cGAS-STING—are known to be deregulated in cancer cells and likely play a role in ISG induction (Dvorkin et al., 2024; Lanng et al., 2024). The cGAS-STING pathway was nevertheless found to be functional in both ALT+ sarcoma cell lines, as STING depletion led to a reduction in ISG transcript levels (Fig 3G and Fig S3F). This observation contrasts with a previous finding that ATRX was necessary for STING activity in ALT+ cells (Chen et al., 2017), as the LB188 and LB857 cell lines lacked ATRX (Fig S3G).

Intriguingly, despite high ISG transcript levels in MG63/TEL+ cells, lysosomal degradation of STING was not increased, while it was clearly detected in LB188/ALT+ sarcoma cells (Fig 3H-I and Fig S3H). Since lysosomal degradation of STING was not detected in LB857/ALT+ sarcoma cells, we reasoned that, in line with our findings in the hybrid cell lines, the extent of STING protein degradation may be linked to CC levels in ALT+ cells. Consistent with this hypothesis, CC levels were significantly higher in LB188/ALT+ than in LB857/ALT+ cells (Fig 3J).

Collectively, these data indicate that a functional cGAS-STING pathway is not incompatible with ALT activity and that the extent of STING lysosomal degradation may be linked to the levels of CC production in ALT+ cells.

### Lysosomes and STING clear ECTRs from the cytoplasm of ALT+ cells

To activate the cGAS-STING pathway in ALT+ cells, ECTRs must be recognized in the cytoplasm. However, using Telo-FISH, we did not detect significant differences in the number of cytosolic ECTRs (cECTRs) between TEL+ and ALT+ cells (Fig 4A-B). It is plausible that cECTRs in TEL+ cells arise from the natural trimming process that occurs at telomeres (Pickett et al., 2009 and 2011; Rivera et al., 2017).

**Figure 4.**
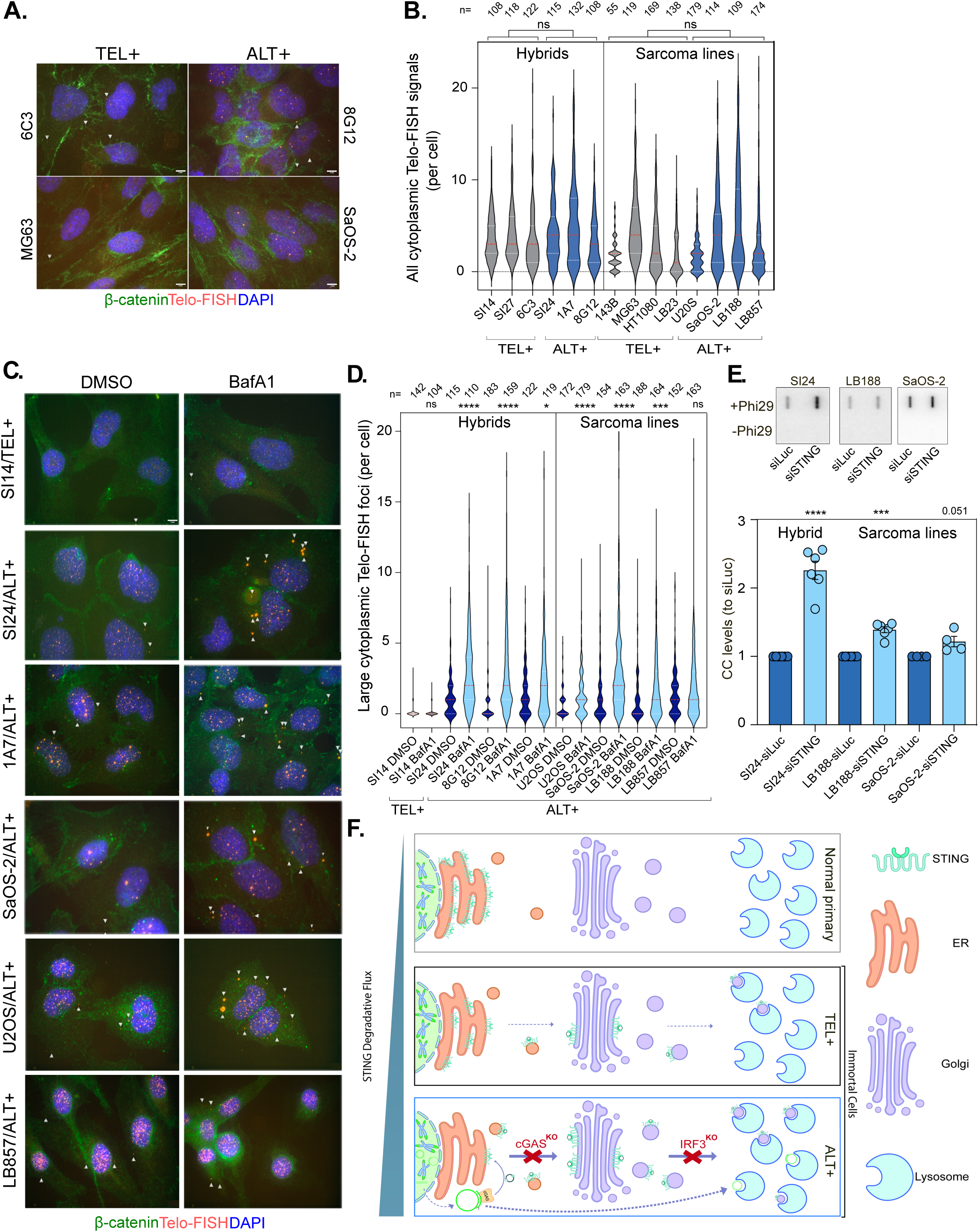
Lysosomes and STING clear ECTRs from the cytoplasm of ALT+ cells. **(A)** Representative pictures of combined immunofluorescence and FISH procedures to stain b-catenin (green), telomeric DNA (red) and DNA (blue) in the indicated cell lines. Scale bar, 5 μm. **(B)** Quantification of cytoplasmic telomeric FISH signals in hybrids (left) and sarcoma cell lines (right). All Telo-FISH signals above background were counted. Violin plots with median (red) and quartiles (white). n=2-3, with the total number of cells counted indicated above. Mann-Whitney tests: TEL+ hybrids or sarcoma cell lines were compared to ALT+ hybrids or sarcoma cell lines. **(C)** Same as A for the indicated cell lines treated (BafA1) or not (DMSO) with 10 nM Bafilomycin A1 for 24 h. n=3. **(D)** Quantification of C. The threshold was increased compared to (B) to monitor the larger Telo-FISH signals. Violin plots with median (red) and quartiles (white). n=3, with the total number of cells counted indicated above. Mann-Whitney tests. **(E)** CC assay on DNA extracted from the indicated cell lines treated with either siLuc or siSTING for 72h. Controls without Phi29 are shown. Mean ± SEM. n=4-6. Ratio paired *t* tests: siSTING samples were compared to siLuc. **(F)** Model. See text for details. For all panels: ns = not significant; \**p* < 0.05; \*\**p* < 0.01; \*\*\**p* < 0.001.

Consistent with the role of cGAS-STING in clearing cytoplasmic DNA *via* lysosomal degradation (Gui et al., 2019), we hypothesized that cECTRs might be degraded as part of the innate immune response. To test this, we first treated cells with BafA1 and analyzed cytoplasmic telomeric signals using FISH. As predicted, treatment with BafA1 led to the appearance of large, bright telomeric DNA foci in the cytoplasm of ALT+ cells, whereas no such signal was observed in TEL+ cells (Fig 4C-D). We then investigated how STING knockdown affects ECTR accumulation. Because transfection reagents interfere with cytoplasmic Telo-FISH assays, we opted to use the CC assay instead. Although this assay does not differentiate between nuclear and cytoplasmic ECTRs—amplifying both DNA species—it still proved useful. This limitation likely explains why the increase in CC levels observed in BafA1-treated SaOS-2/ALT+ cells appeared less pronounced compared to the cECTR-specific signal detected by FISH (Fig S4A-B). Nonetheless, after validating the CC assay’s ability to detect cECTR accumulation, we depleted STING in SaOS-2/ALT+, SI24/ALT+, and LB188/ALT+ cells. This led to a significant rise in CC levels across all three lines (Fig 4E and Fig S4C), reinforcing the idea that STING plays a key role in clearing ECTRs from the cytoplasm of ALT+ cells.

Collectively, these results demonstrate the role of lysosomes and STING in clearing ECTRs from the cytoplasm of ALT+ cells.

## Discussion

Initially recognized as a crucial signaling pathway for detecting pathogenic DNA and triggering an innate immune response *via* the expression of interferon-sensitive genes, it has since been established that cGAS can also be activated by endogenous self-DNA species, including extranuclear chromatin released in response to genotoxic stress or DNA fragments that escape from mitochondria (Hopfner and Hornung, 2020). Likewise, ECTRs, when induced experimentally by tethering nucleases to telomeres, are potent activators of the cGAS-STING pathway (Chen et al., 2017). In line with this, ALT+ cells, which naturally and chronically produce ECTRs, have been suggested to require *STING* gene silencing to prevent the harmful effects of continuous cGAS-STING pathway activation (Chen et al., 2017). Lysosomal degradation of STING is yet another defense mechanism that prevents the detrimental effects of prolonged ISG induction (Ahn et al., 2012). Our findings indicate that the frequent loss of STING protein in ALT+ cells is primarily due to its lysosomal degradation (Fig 4F), thus refining the model of immune sensing in ALT+ cells.

Our findings support the notion of chronic activation of the cGAS-STING pathway in ALT+ cells, as 2’3’-cGAMP levels were significantly higher in ALT+ cells compared to TEL+ and primary human cells, and ISG expression was notably elevated in ALT+ cells relative to TEL+ isogenic hybrid cell lines. The correlation between ISG expression and CC levels in ALT+ hybrids suggests that cytoplasmic ECTRs may play a role in activating the cGAS-STING pathway, leading to the subsequent lysosomal degradation of STING. In line with this, cGAS deletion reduced ISG expression and restored STING protein levels. Moreover, while we observed cGAS- and lysosomal-dependent STING degradation, our finding that IRF3, but not TBK1, is essential for STING’s lysosomal degradation suggests that this process occurs independently of IRF3’s transcriptional activity and the induction of ISGs which typically requires IRF3 phosphorylation by TBK1. This also reveals a novel, TBK1-independent function of IRF3 in modulating the innate immune response. We propose that IRF3 when bound to STING, may act as an adaptor that directs STING for degradation. Although TBK1 has been identified as the kinase responsible for phosphorylating the S366 residue of STING, facilitating its interaction with IRF3 at the Golgi (Liu et al., 2015; Jeltema et al., 2023), other studies suggest that the ULK1 kinase may also contribute to this process (Konno et al., 2013). In line with the notion that STING degradation is independent of an increased macroautophagic flux and does not rely on ATG5 or LC3 lipidation (Gui et al., 2019), we observed no increase in the LC3-II/LC3-I ratio in ALT+ hybrid cells with enhanced STING degradation. Similarly, RNA-seq analysis did not show any consistent upregulation of TFEB-regulated genes (Sardiello et al., 2009) in ALT+ hybrids (Fig S4D), indicating that the cGAS-STING-TFEB axis, which mediates lysosomal biogenesis (Lv et al., 2024, Huang et al., 2025; Tang et al., 2025; Xu et al., 2025; Tapia et al., 2025), was not hyper-activated in ALT+ cells.

Why do most ALT+ cells continue to express cGAS and STING instead of silencing these genes, which would effectively prevent the harmful activation of ISGs? First, our findings suggest that STING has an active role in promoting the lysosomal degradation of cECTRs in ALT+ cells. By clearing cytosolic DNA and reducing the release of telomeric DNA from tumor cells into the microenvironment, this process limits the activation of immune cells, such as dendritic cells, thereby providing an advantage to cancer cells (Xu et al., 2017; Mender et al., 2020). We propose that ECTRs are initially more prevalent in the cytoplasm of ALT+ cells but are later degraded in the lysosome. Considering the absence of evidence for a significant increase in macroautophagy in ALT+ cells, this suggests the possibility that a microautophagy process could be involved, potentially through the DNautophagy process, which involves a direct interaction between the LAMP2C lysosomal membrane protein and DNA (Fujiwara et al., 2013). Our finding that ECTRs were also present in the cytoplasm of TEL+ cells, without a corresponding increase in their abundance upon bafilomycin A1 treatment, leads us to hypothesize that the cECTRs in ALT+ cells could be longer or structurally different from those in TEL+ cells. This difference may make them more effective in inducing their degradation and potentially STING degradation. In this context, it’s worth noting that cGAS preferentially recognizes longer DNA molecules (Luecke et al., 2017).

Secondly, we propose that maintaining an active cGAS-STING pathway in ALT+ cells may support their survival by promoting pro-tumoral activity through NF-κB and IL-6, as seen in cancer cells with high chromosomal instability (Bakhoum et al., 2018; Hong et al., 2022; Lanng et al., 2024). Downstream of the IL-6/JAK signaling pathway, ERK1/2 plays a pivotal role in cancer progression by regulating processes such as proliferation, migration, and stress responses (Guo et al., 2020), which could provide a survival advantage to ALT+ cancer cells. Supporting this, both NF-κB and IL-6 signaling were significantly upregulated in ALT+ cells relative to TEL+ hybrids (Fig S4E-F). This opens interesting therapeutic possibilities, and the proposed strategy of combining ERK1/2 inhibitors with STING agonists as boosters of anti-tumor immunity (Lanng et al., 2024) may be applicable to ALT+ tumors. It is, however, possible that the chronic cGAS-STING activation in ALT+ cells may be sufficient, making the addition of STING agonists unnecessary and potentially reducing the harmful effects associated with STING agonist-induced immune cell death (Sivick et al., 2018).

Collectively, our findings expand our understanding of immune sensing in ALT+ cancers by demonstrating that complete loss of cGAS or STING is not necessary for ALT emergence, as many ALT+ cancer cells maintain a functional cGAS-STING axis. This underscores promising therapeutic possibilities for cancer immunotherapy, especially if chronic cGAS-STING activation in ALT+ cells enhances the recruitment of cytotoxic immune cells to the tumor microenvironment, provided that the downstream pro-tumoral effects of STING are inhibited.

## Supporting information

Key Resources table

## Acknowledgements

We apologize to our colleagues whose excellent work could not be cited due to space limitations. A.D., M.V. and M.M. are supported by the Fonds National de la Recherche Scientifique (FNRS) and UCLouvain. D.G. and A.D. are supported by the Télévie/FNRS. A.L. is supported by the de Duve Institute. J.N. is supported by the National Cancer Institute (K99CA252447 and R00CA252447) and Cancer League of Colorado. R.H. is supported by the National Cancer Institute (T32CA190216-08). We are grateful to the de Duve Institute for constant support. We are grateful to Jan Karlseder and Roddy O’Sullivan for stimulating discussions and insightful suggestions.

## Materials and Methods

Detailed references of reagents are provided in the Key Resources Table.

### Cell culture and cell treatments

For Fig 1 and Fig 2, all cells were grown at 37 °C under 7.5% CO_2_ and 3% O_2_ and all media were supplemented with 0.1 mM non-essential amino acids (NEAA) (Corning), 15% fetal bovine serum (FBS) (VWR), 100 IU/mL Penicillin and 100 μg/ml Streptomycin (Gibco). IMR90, MRC5, WI38, HeLa-LT, KMST-6, WI38-VA13, GM847, Calu-3, HOS, SaOS-2, and SK-N-F1 cells were grown in GlutaMax-DMEM (Gibco); U2OS, HT-29 and G292 cells were grown in McCoy’s 5A (modified) media (Fisher Scientific); SJCRH30 cells were grown in RPMI-1640 (Fisher Scientific) and TOV-112D cells were grown in a 1:1 mixture Medium 199 (Sigma-Aldrich) with a final concentration of 1.5 g/L sodium bicarbonate (Sigma-Aldrich), and MCDB 105 (Cell Applications) with a final concentration of 2.2 g/L sodium bicarbonate. For Fig 3 and Fig 4, cells were grown at 37°C under 5% CO_2_ and all media were supplemented with 10% FBS (Cytiva), 1% NEAA (Gibco) and 1% penicillin/streptomycin (Capricorn Scientific). SI14, SI27, 6C3, SI24, 1A7 and 8G12 hybrids (Episkopou et al., 2014 and 2019; Raghunandan et al., 2021) were grown in EMEM (Gibco); 143B, MG63, HT1080, U2OS and SaOS-2 cells were grown in DMEM (Gibco); LB23, LB188 and LB857 cells were cultured in IMDM (Gibco). All cells have been tested to be free of mycoplasma.

Cells were treated with drugs as follows: 50 nM Bafilomycin A1 (BafA1) (Tocris) (Fig 1B and Fig S1D) or 10 nM BafA1 (Sigma-Aldrich) (Fig S3C and S3H, Fig 4C-D and Fig S4A-B), 50 µM hydroxychloroquine (HCQ) (Tocris), or 1 µM MG132 (R&D Systems) for 24 h. Cells were treated with herring testes DNA (HT-DNA) (Sigma-Aldrich) at 2 µg/mL, poly(dA:dT) (InvivoGen) at 2 µg/mL, or 2’3’-cGAMP (InvivoGen) at 10 µg/mL for indicated time points.

### Gene knockout

To generate pooled knockout cell populations, lentiCRISPRv2 was used. The plasmid was obtained from F. Zhang through Addgene (plasmid 52961) (Sanjana et al., 2014) and the guide target sequence was cloned as phosphorylated adapter in the *Bsm*BI-digested lentiCRISPRv2 vector using T4 PNK and T4 ligase (NEB). The guide RNA sequences were as follows: sgGFP 5′-GAAGTTCGAGGGCGACACCC-(PAM)-3′, sgIRF3 A 5’-TTGGAAGCACGGCCTACGGC-(PAM)-3’, sgIRF3 B 5’-CGAGCCTCTTGGTCCACGGC-(PAM)-3’, sgTBK1 A 5’-TCCACGTTATGATTTAGACG-(PAM)-3’, sgTBK1 B 5’-TAATGCGAAAGGGGATACGA-(PAM)-3’, sgcGAS A 5’-GCTTCCGCACGGAATGCCAG-(PAM)-3’, and sgcGAS B 5’-AGACTCGGTGGGATCCATCG-(PAM)-3’ (Integrated DNA Technologies). Production of lentivirus was performed as described previously (Arnoult et al., 2017). In brief, HEK293T cells were transfected with 7 µg of DNA using Lenti-X Packaging Single-Shot system (Takara Bio). Then, 48 h after transfection, viral supernatant was collected, mixed with Lenti-X-Concentrator (Takara Bio), and incubated at 4 °C overnight. Following incubation, the supernatant was centrifuged, resuspended in media supplemented with serum and used for transduction in the presence of Lenti-Blast (Oz Biosciences). After transduction, cells were selected with 1 μg/mL puromycin (Thermo Scientific) for 2 days and KO cell populations were tested by western blotting for the absence of protein staining.

### Knockdown with siRNAs

Transfections with 100 nM siRNAs (listed in the Key Resources Table) were performed using Genius DNA Transfection reagent (Westburg) (Fig 3B and 3G) or Lipofectamine 2000 (Invitrogen) (Fig 4E, S4A-C) according to the manufacturer’s instructions. Cells were collected 72 h post-transfection. For Fig S4 A-B, 48 h post-transfection, medium was replaced by medium containing 10 nM Bafilomycin A1 and incubated for a further 24 h at 37°C with 5% CO_2_. For Fig S2, 60 nM ON-TARGETplus SMART pools (Horizon Discovery, listed in the Key Resources Table) were transfected using the Lipofectamine RNAiMAX kit (Thermo Fisher Scientific) according to the manufacturer’s instructions and transfections were repeated twice, 24 h apart, before cell collection 72 h after the second transfection.

### RNA extraction, cDNA synthesis and qPCR

RNA was extracted using RNeasy Mini Kit (Qiagen) according to the manufacturer’s instructions. Genomic DNA was eliminated by double digestion with DNase I (Qiagen). For Fig 3B, 3D, 3F, 3G, 3I, S3A, S3B and S3E, cDNA was synthesized using 1 µg RNA, 200 U of M-MLV RT (Thermo Fisher Scientific), 0.5 mM dNTPs (Thermo Fisher Scientific), 10 mM DTT (Thermo Fisher Scientific), 10 ng/µL random hexamers (Thermo Fisher Scientific) and 20 U of Ribolock RNAse inhibitor (Thermo Fisher Scientific). Samples were incubated at 42°C for 1 h before inactivation at 95°C for 5 min. qPCRs were performed using KAPA SYBR FAST (Sigma-Aldrich) and primers listed in the Key Resources Table. For Fig S1C-D, cDNA was synthesized from 3 µg RNA using the SuperScript III First-Strand Synthesis System (Thermo Fisher Scientific) and qPCRs were performed using Power SYBR Green Master Mix (Thermo Fisher Scientific) and primers listed in the Key Resources Table. In all cases, relative gene expression was normalized using *ACTB* as a housekeeping gene and calculated using the comparative Ct (ΔΔCT) method.

### Western Blotting

Immunoblots of Fig 1A-B, 2A, 2C, S1A and S2 were performed as described previously (O’Sullivan et al., 2010). In brief, cells were lysed in lysis buffer containing 211 mM Tris-HCl, 280 mM Tris base, 147 mM LDS and 2.17 M glycerol. Samples were normalized by protein concentration using the DeNovix DS-11 FX+. Proteins were resolved using NuPage Bis-Tris gel electrophoresis (Invitrogen, NP0342, NP0321 or WG1402) and transferred to nitrocellulose membranes (Bio-Rad). Primary antibodies, along with the dilutions used, are listed in the Key Resources Table. Secondary antibody was peroxidase-conjugated anti-Rabbit IgG (Cell Signaling Technologies). Peroxidase activity was detected using an ECL kit (Genesee Scientific) and the Bio-Rad ChemiDoc imager. Immunoblots of Fig 3B, 3C, 3H, S3C-D, S3F-H and S4C, cell extracts were prepared with RIPA buffer (50 mM Tris-HCl pH 7.4, 150 mM NaCl, 1 % NP40, 0.25 % sodium deoxycholate) supplemented with protease inhibitors and phosphatase inhibitors. Western blotting was performed according to standard procedures using 10-20 µg of proteins. PVDF membranes were blocked with either 5% powdered milk or BSA for 1 h at room temperature. Primary antibodies, along with the dilutions used, are listed in the Key Resources Table. Secondary antibodies were goat peroxidase-conjugated anti-Rabbit IgG (Enzo Life Sciences) or anti-Mouse IgG (Abcam). SuperSignal West Pico Chemiluminescent Substrate was used for revelation and signals were quantified using ImageJ software.

### cGAMP ELISA

Cells were lysed in M-PER lysis buffer (Thermo Fisher Scientific) according to the manufacturer’s instructions. Samples were normalized by protein concentration using the DeNovix DS-11 FX+. Intracellular 2’3’-cGAMP concentrations were measured using the 2’3’-cGAMP ELISA Kit (VWR) according to the manufacturer’s instructions.

### Immunofluorescence and telomeric FISH

Primary and secondary antibodies, along with the dilutions used, are listed in the Key Resources Table. For IF experiments, cells were seeded onto poly-L-Lysine-coated glass coverslips. For Fig 1C and S1E, after 24 h incubation on coverslips, cells were fixed in 4% paraformaldehyde (PFA) in PBS for 10 min, washed in PBS and incubated in blocking solution of 5% BSA in PBS at room temperature. Cells were then incubated with primary antibodies (anti-LAMP2 (2D3B9, Protein Tech), anti-calnexin (AF18, Invitrogen) or anti-TMEM173/STING (Protein Tech) for 2 h, washed in PBS and incubated with secondary antibodies: goat anti-Rabbit IgG (Thermo Fisher Scientific) or donkey anti-Mouse IgG (Thermo Fisher Scientific) for 1 h at room temperature. The samples were finally washed in PBS and mounted in ProLong Diamond with DAPI (Invitrogen). Imaging was performed using the Zeiss Axio Imager.Z2 and ImageJ was used for image analysis. For telomeric DNA FISH after IF (Fig 4A and 4C), cells were incubated overnight on coverslips before processing. For Fig 4C, after overnight incubation of the cells on coverslips, medium was replaced by fresh medium containing either 10 nM BafA1 or DMSO and cells were incubated for 24 h before processing. After incubation, medium was removed, coverslips were rinsed with PBS, fixed with 4% PFA in PBS for 15 min, washed in PBS three times and incubated in permeabilization buffer (20 mM Tris-HCl pH 8, 50 mM NaCl, 3 mM MgCl_2_, 300 mM sucrose, 0.5% Triton X-100) for 10 min. After a new wash in PBS, cells were incubated in blocking solution (3% BSA, 0.1% Triton X-100 in PBS) for 1 h at 37°C, before overnight incubation at 4°C with anti-β-Catenin antibody (D10A8, Cell Signaling Technology, 1:200) diluted in blocking solution. After three washes of 3 min each at 45°C with 0.1% Tween in PBS, coverslips were incubated with goat anti-Rabbit IgG (Alexa Fluor™ 488 conjugated, 1:400) (Invitrogen) for 40 min at 37°C. Coverslips were then washed three times with 0.1% Tween in PBS at 45°C and three times with PBS at room temperature before a second fixation with 4% PFA for 2 min. Coverslips were then rinsed with PBS and incubated with 0.1 mg/mL RNase A, diluted in 2x SSC, for 1 h at 37°C. After three washes with 2x SSC and one with PBS, coverslips were fixed again with 4% PFA for 2 min before dehydration through serial incubations of 2 min each in 70%, 80%, 90% and 100% ethanol before air-drying. Coverslips were then incubated with 160 nM (TYE563)GGGTtAGGGttAGgGTTAGGGttAGGGttAGGGtTA(TYE563) TeloG LNA^TM^ probe (Qiagen, with small letters indicating LNA^TM^ modified bases) in hybridisation solution (50% (v:v) deionised formamide, 2x SSC, 1x Blocking Reagent) before incubation for 3 min at 83°C. Unbound probe was washed off twice with formamide solution (50% formamide, 20 mM Tris, 2x SCC) for 15 min and three times with Tris solution (50 mM Tris pH 7.4, 150 mM NaCl, 0.05% Tween 20) for 5 min. Coverslips were serially dehydrated again, air dried and mounted with Fluorescence Mounting Medium (Agilent) containing 1 μg/mL DAPI. Images were acquired with the Cell Observer Spinning Disk confocal microscope (Zeiss) with 100x oil objective and analysed using Fiji software. Counting was done as followed: after ZProjection at Max intensity, the same thresholds were applied for all samples, and FISH signals in cytoplasm were counted manually.

### RNA-seq analysis of TEL^+^ and ALT^+^ SV40 fibroblasts

RNA was extracted using RNeasy Mini Kit (Qiagen) according to the manufacturer’s instructions and genomic DNA was eliminated by double digestion with DNase I (Qiagen). RNA samples were sent to Macrogen for transcriptome sequencing. Fastq files were processed using a standard RNA-seq pipeline including Trimmomatic-0.38 (Bolger et al., 2014) to remove low quality reads, hisat2-2.1.0 (Kim et al., 2015) to align reads to the human genome (GRCh38) and gene expression levels were evaluated using featureCounts from subread-2.0.0 (Liao et al., 2014) and Homo_sapiens.GRCh38.94.gtf. Differential expression analyses were performed with DESeq2 Bioconductor package v1.46.0 (Love et al., 2014). Gene set enrichment analyses were done using clusterProfiler v4.14.4 (Yu et al., 2012).

### C-circle assay

The protocol established by Henson et al. (2017) was adapted as follows. Cells were resuspended in lysis buffer (10 mM Tris-HCl pH 8, 1 mM EDTA, 0.5% SDS, 10 mM NaCl) supplemented with 50 μg/mL RNase A (Thermo Fisher Scientific) and incubated for at least 1 h at 37°C. Cell lysate was supplemented with 100 μg/mL proteinase K (Sigma-Aldrich) and incubated overnight at 37°C. DNA was purified with phenol-chloroform-isoamyl alcohol (25:24:1) followed by isopropanol precipitation in 0.3 M sodium acetate, pH 5.5. Two to five µg of genomic DNA were digested with 20 U of *Rsa*I/*Hinf*I (NEB), and purified with phenol-chloroform-isoamyl alcohol (25:24:1) followed by isopropanol precipitation in 0.3 M sodium acetate, pH 5.5. Finally, 30 ng of digested genomic DNA were incubated with or without 7.5 U of Phi29 DNA polymerase in reaction mix (4 μg/mL BSA, 4 mM DTT, 0.1% Tween-20, 1 mM each dATP, dCTP, dGTP and dTTP, 1X Phi29 Buffer) at 30°C for 8 h and then at 65°C for 20 min. Amplification products were slot-blotted onto Hybond N^+^ nylon membrane (Cytiva) before UV-crosslinking with 1200 µJ (UV Stratalinker™ 1800, Stratagene). Next, the membrane was pre-hybridized for 1 h at 42°C in ULTRAhyb™-Oligo solution (Invitrogen), before overnight incubation at 42°C in hybridization solution containing the radioactive telomeric probe prepared as follows: the (*CCCTAA*)_4_ Telo-C oligonucleotide at 10 μM was denatured for 5 min at 68°C. One μL of the probe was then radioactively labeled with 6 μL [γ-32^P^]ATP (10 mCi/mL) (Revvity) and 1 μL of T4 PNK (10 U/μl) (Roche) in a total volume of 20 μL of PNK buffer 1x. The final mix was incubated for 20 min at 37°C before addition of 2 μL EDTA 0.5 M pH 8.0. Forty μl of 1x PNK buffer were then added to the radioactive probe before purification on a G-25 column (Cytiva). After hybridization, the radioactive solution was discarded and the membrane was washed once for 10 min at room temperature with stringent solution I (2X SSC, 0.1% SDS), once for 3 min at 42°C with stringent solution II (0.2X SSC, 0.1% SDS) and twice for 5 min at room temperature with stringent solution I. Finally, a Phosphorscreen was exposed to the membrane, and read on the Typhoon Trio Phosphor Imager.

### Statistical analyses

Graphpad Prism version 10 was used to perform the statistical analyses: all datasets were checked for the normality of their distribution with the Shapiro-Wilk test. The statistical tests that were used are detailed in the figure legends.

