## Supplementary figures and images for "Constitutive cGAS-STING activation in ALT+ cells"

### Fig S1

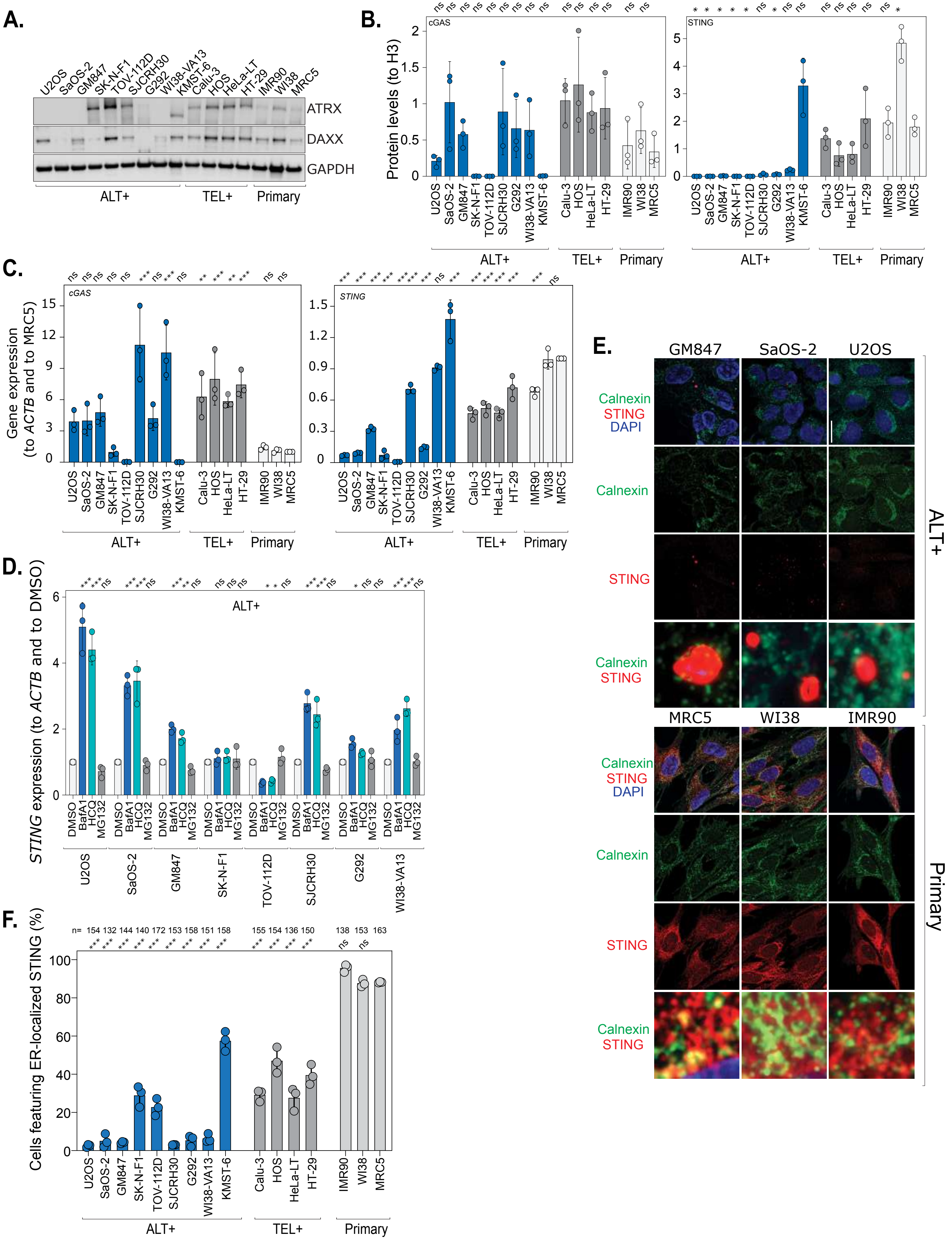

### Fig S2

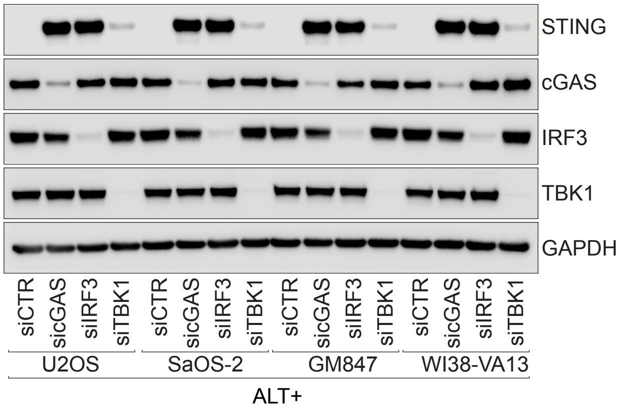

### Fig S3

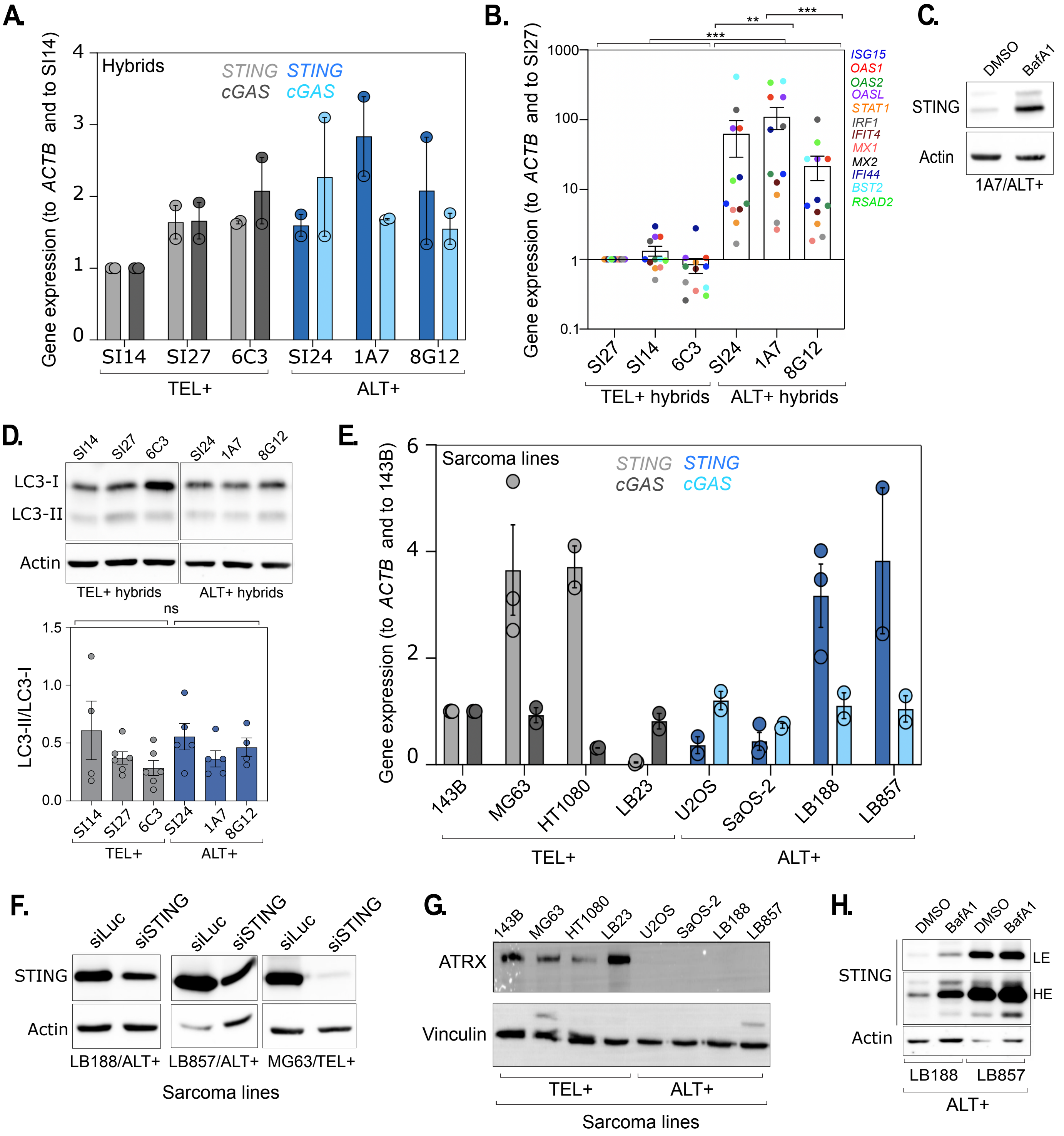

### Fig S4

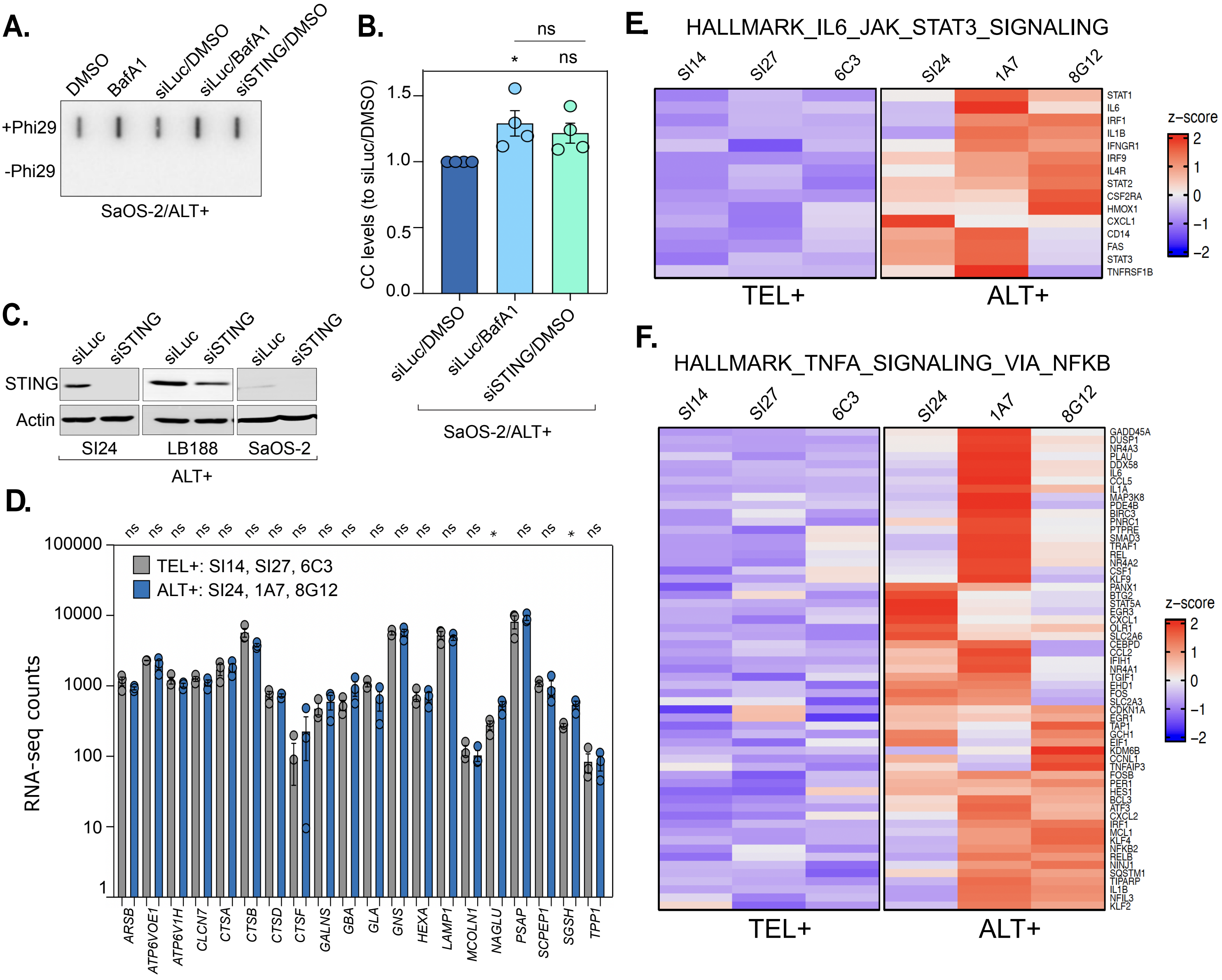
