## Supplementary material for "Constitutive cGAS-STING activation in ALT+ cells": Suppl legends

**Figure S1. Expanded data for Fig 1. (A)** Immunoblot analysis of ATRX and DAXX in primary human fibroblasts (IMR90, WI38, MRC5), ALT+ cells (U2OS, SaOS-2, GM847, SK-N-F1, TOV-112D, SJCRH30, G292, WI38-VA13, KMST-6), and TEL+ cancer cells (Calu-3, HOS, HeLa-LT, HT-29). GAPDH was used as a loading control. n=3. **(B)** Quantifications of Fig 1A. cGAS (left) and STING (right) protein levels were normalized to H3. Mean ± SD. n=3. Brown-Forsythe ANOVA test: all samples were compared individually to primary MRC5 cells. **(C)** RT-qPCR analysis of *cGAS* (left) and *STING* (right) mRNA levels in the indicated primary, ALT+, or TEL+ cells, normalized to *ACTB* and to MRC5. Mean ± SD. n=3. Brown-Forsythe and Welch ANOVA test: all samples were compared individually to primary MRC5 cells. **(D)** RT-qPCR analysis of *STING* mRNA levels in the indicated ALT+ cells treated for 24 h with either DMSO (control), the lysosomal inhibitors bafilomycin A1 (BafA1) or hydroxychloroquine (HCQ), or the proteasome inhibitor MG132. Levels were normalized to *ACTB* first and then to DMSO control. Mean ± SD. n=3. Two-way ANOVA test: all samples were compared individually to DMSO-treated cells. **(E)** Apotome maximum intensity projection images of ALT+ cells (U2OS, SaOS-2, GM847) or primary human fibroblasts (IMR90, WI38, MRC5), co-immunostained with calnexin and STING. n=3. Scale bar, 20 µm. **(F)** Quantifications of (E). Percentage of cells with ER-localized STING. Mean ± SD. n=3, with the total number of cells counted indicated above. Brown-Forsythe and Welch ANOVA test: all samples were compared individually to primary MRC5 cells.
For all panels: ns = not significant; **p* < 0.05; ***p* < 0.01; ****p* < 0.001.

**Figure S2. Expanded data for Fig 2.** Immunoblot analysis of ALT+ cells transfected with either non-targeting siRNA (siCTR) or siRNA targeting cGAS, IRF3, or TBK1. Protein extracts were collected 72 h post-transfection. GAPDH was used as a loading control. n=3.

**Figure S3. Expanded data for Fig 3. (A)** RT-qPCR analysis of *cGAS* and *STING* in the hybrid cell lines, normalized first to *ACTB* expression levels and then to SI14. Mean ± SEM. n=2. **(B)** Validation of RNA-seq by RT-qPCR on selected ISGs in the hybrid cell lines, normalized first to *ACTB* expression levels and then to SI27. Mean ± SEM. n=1-3. Two-tailed Mann-Whitney tests. ***p* < 0.01; ****p* < 0.001. **(C)** Recovery of STING protein in 1A7/ALT+ cells treated with 10 nM Bafilomycin A1 (BafA1) for 24 h. n=2. **(D)** Above: representative immunoblot analysis of LC3-I and LC3-II in the hybrids. Below: quantification of LC3-II/LC3-I ratios. Mean ± SEM. n=4. **(E)** RT-qPCR analysis of *cGAS* and *STING* in the sarcoma cell lines, normalized first to *ACTB* expression levels and then to 143B. Mean ± SEM. n=2-3. **(F)** Representative immunoblot analyses of STING knock-down in the indicated sarcoma cell lines for Fig 3G. Actin was used as a loading control. **(G)** Immunoblot analysis of ATRX in the indicated sarcoma cell lines. Vinculin was used as a loading control. **(H)** Same as (C) in LB188/ALT+ and LB857/ALT+ sarcoma cell lines. n=3. LE (low exposure), HE (high exposure). Actin was used as a loading control.

**Figure S4. Expanded data for Fig 4 and Discussion. (A)** CC assay on DNA extracted from the indicated cell lines treated with either siLuc or siSTING for 72h. When indicated, cells were treated with 10 mM Bafilomycin A1 (BafA1) or control DMSO for the last 24 h. Controls without Phi29 are shown. **(B)** Quantifications of (A). Values were normalized to siLuc/DMSO. Mean ± SEM. n=4. Ratio paired *t* tests. **(C)** Representative immunoblot analyses of STING knock-down in the indicated cell lines for Fig 4E. Actin was used as a loading control. **(D)** Counts for the indicated transcripts obtained by RNA-seq in the TEL+ (SI14, SI27, 6C3) and ALT+ (SI24, 1A7, 8G12) hybrids. Mean ± SEM. Unpaired *t* tests. **(E-F)** Heatmaps from RNA-seq analysis in TEL+ and ALT+ hybrids: Hallmark_IL6_JAK_STAT3 signaling (E) or HALLMARK_TNFA_SIGNALING_VIA_NFKB (F).

For all panels: ns = not significant; *$p < 0.05$.
